# Delaying cefiderocol resistance development in NDM-producing *Enterobacter cloacae* complex by combining cefiderocol with aztreonam *in vitro*

**DOI:** 10.1101/2024.02.13.579981

**Authors:** Lisa Göpel, Minh Thu Tran Nguyen, Tung Tran Thanh, Susanne Hauswaldt, Özge Nur Canbulat, Jan Rupp, Sébastien Boutin, Dennis Nurjadi

## Abstract

**Background:** The rapid development of cefiderocol resistance poses a significant concern, particularly in Enterobacterales that produce New Delhi metallo-β-lactamase (NDM). This study explores the potential of inhibiting the development of cefiderocol resistance by combining cefiderocol with aztreonam.

**Methods:** A resistance induction experiment using 20 clinical isolates was performed to assess the impact of cefiderocol-aztreonam on preventing cefiderocol resistance development at 4x and 10x cefiderocol MIC, with and without aztreonam (2, 4, 8 µg/ml). Additionally, serial passaging with doubling cefiderocol concentrations was performed with and without aztreonam. Whole genome sequencing (WGS) was performed to identify potential genetic factors associated with the phenotype.

**Results:** Among the 20 *E. cloacae* complex isolates, 40% (8/20) exhibited a significant reduction in cefiderocol MIC (≥4-fold MIC reduction) in the presence of 4 µg/ml aztreonam. Combining cefiderocol with a fixed concentration of 4 µg/ml aztreonam inhibited cefiderocol resistance development in these eight isolates at an inoculum of 10^7^ cfu/ml. Additional resistance induction experiments through serial passaging indicated a delayed emergence of cefiderocol-resistant clones when cefiderocol was combined with aztreonam. WGS analysis revealed a significant positive association between *bla*_CTX-M-15_, *bla*_OXA-1_, and other co-localized genes with a substantial MIC reduction for cefiderocol-aztreonam compared to cefiderocol alone.

**Conclusion:** Our study suggested that cefiderocol resistance development in NDM-producing *E. cloacae* complex can be delayed or inhibited by combining cefiderocol with aztreonam, even in the presence of multiple β-lactamase genes. A MIC reduction of at least 4-fold emerges as the most reliable predictor for inhibiting resistance development with this dual β-lactam combination.

## Background

Cefiderocol, a relatively novel siderophore-conjugated cephalosporin, is a promising agent to treat infections caused by carbapenem-resistant Enterobacterales, including those producing the New Delhi metallo-β-lactamase (NDM). However, minimum inhibitory concentrations (MIC) of cefiderocol in NDM-producing Enterobacterales are generally higher than in strains carrying other β-lactamases [1].

In a prior study, we identified that NDM-production facilitates cefiderocol resistance development in *Enterobacter cloacae* complex, and the resistance development could be prevented by inhibiting NDM activity with dipicolinic acid in vitro [2]. However, dipicolinic acid is toxic and not approved for clinical use. Therefore, other strategies to inhibit or bypass NDM activity by combining clinically approved drugs may be an option.

The combination of double β-lactam antibiotics to overcome this challenge has not been studied extensively. NDM-type metallo-β-lactamases can hydrolyze penicillins, cephalosporins, and carbapenems. On the other hand, the monobactam aztreonam is not susceptible to NDM hydrolysis [3, 4]. However, aztreonam can be hydrolyzed by extended-spectrum β-lactamases, AmpC β-lactamases, or OXA-48-like carbapenemases, which are often co-produced by NDM-producing Enterobacterales [5].

Given the stability of cefiderocol against NDM and other beta-lactamases and the stability of aztreonam against NDM, we hypothesize that the combination of aztreonam and cefiderocol may result in “cross-protection” effect of both antibiotic substances and may be capable of preventing NDM-mediated development of cefiderocol resistance, making this combination an interesting and promising candidate for alternative therapeutic strategies against NDM-producing carbapenem-resistant Enterobacterales.

## Methods

### Cefiderocol susceptibility testing

Antibiotic susceptibility testing (AST) for cefiderocol was performed using disk diffusion and broth microdilution according to EUCAST recommendations. Disk diffusion was performed using cefiderocol (30µg, Mast Group, Germany) and aztreonam disks (30µg, Mast group, Germany). Broth microdilution for aztreonam (Supelco, USA) was performed using cation-adjusted Mueller-Hinton broth (CA-MHB) (Sigma-Aldrich, Germany) and for cefiderocol (Shionogi, Japan) with iron-depleted CA-MHB, which was prepared according to a published protocol [6]. *Escherichia coli* ATCC®25922 was used as a quality control strain. Synergy test was determined by a checkerboard assay (Supplementary methods).

### Phenotypic characterization of clinical isolates

The clinical NDM-producing *E. cloacae* complex isolates were collected from routine diagnostics. Species identification was performed using MALDI-TOF MS (Bruker, Germany). Initial susceptibility testing was performed using Vitek®2 (Biomérieux, Germany) and was re-interpreted using current EUCAST clinical breakpoints for this study (v13.1). The selection criteria for the inclusion of these isolates in this study was NDM-positivity, which was determined by PCR and confirmed by Illumina whole-genome sequencing for this study.

### WGS and bioinformatics analysis

DNA extraction was performed from fresh culture on BD™ Columbia Agar with 5% Sheep Blood (Becton Dickinson GmbH, Heidelberg, Germany) at 37 °C. DNA was extracted using the DNeasy Blood and Tissue Kit (Qiagen GmbH, Hilden, Germany) according to the manufacturer’s protocol. Library preparation was performed using Nextera DNA Flex Library Prep Kit (Illumina) and sequencing was done on a MiSeq Illumina platform (short-read sequencing, 2□×301 bp). Post-sequencing procedure is described in detail as a supplementary method. The gene presence/absence table from Roary was used to associate gene variability with the phenotypic resistance to the combination of aztreonam and cefiderocol using Pyseer v1.3.11 with a fixed model including the recombination adjusted phylogenetic distance and using 2 dimensions based on the MDS eigenvalues (Supplementary Figure S1). Only genes associated with a Likelihood ratio test p-value ≤ 0.01 were considered as significantly associated with the phenotype.

### Resistance induction in vitro using single antibiotic exposure

A 3.0 McFarland suspension (equivalent to approximately 1×10^9^ cfu/ml) was prepared in fresh iron-depleted CA-MHB from a fresh overnight culture, 100µl of the suspension was a defined inoculum (range: 10^4^ – 10^8^ cfu/ml) was transferred to sterile culture tubes containing 5 mL iron-depleted CA-MHB with (i) 4xMIC cefiderocol and (ii) 10xMIC cefiderocol, supplemented with 2, 4 and 8 µg/ml aztreonam for the induction experiment with *E. hormaechei* etcl_1. Bacterial growth was determined after 24 h of incubation (37°C, 200 rpm) by measuring at OD_600_ using standard photometer cuvettes (Sarstedt, Germany). An identical protocol was applied for the 20 clinical NDM-producing isolates with the only modification of using a fixed concentration of 4 µg/ml of aztreonam.

### Resistance induction by serial passaging

100µl from an overnight liquid culture of *E. cloacae* growing in 5 mL iron-depleted CA-MHB was subcultured, on a daily basis, into a 5 mL fresh media containing cefiderocol, aztreonam, cefiderocol-aztreonam and without antibiotics. The concentration of aztreonam was kept constant at 4 µg/ml, while the concentration of cefiderocol was doubled daily (starting from 0.5 µg/ml) until the concentration of 16 µg/ml was reached. Bacterial growth was determined using a photometer (OD600). After each passage, a cefiderocol disk diffusion test from the liquid culture adjusted to 0.5 McFarland in sterile 0.9% NaCl was performed to detect the development of resistance.

### Statistical analysis

Data visualization and descriptive statistical analysis were performed using GraphPad Prism version 9 (GraphPad, USA), Stata18/BE (StataCorp, USA) or R 4.3.0. A p-value of <0.05 was considered as statistically significant.

### Ethical considerations

The usage of clinical bacterial strains without patient data does not require ethical clearance.

## Results

### The combination of cefiderocol and aztreonam can reduce cefiderocol MIC and prevent the emergence of resistance

To investigate the interaction between cefiderocol and aztreonam, we performed a checkerboard assay using an NDM- and OXA-48-expressing *E. hormaechei* (*E. cloacae* complex) etcl_1 from our previous studies [2, 7]. In the presence of 4 µg/ml aztreonam, the cefiderocol MIC was reduced by 16-fold from 8 µg/ml to 0.5 µg/ml. Calculation of the fractional inhibitory concentration index suggested an additive effect but no synergistic effect (FICI=0.5). For a cefiderocol-resistant mutant of etcl_1 (etcl_1_mut) due to a frameshift mutation in the *cirA* gene, combining cefiderocol with aztreonam did not exhibit any significant effect (Table 1, Supplementary Figure S2).

**Table 1.** Cefiderocol and aztreonam susceptibility profiles of *Enterobacter cloacae* complex isolates used in this study.

| ID | Species (WGS) | MLST<sup>a</sup> | carbapenemase | CTX-M<sup>b</sup> | Cefiderocol BMD<sup>c,d</sup> |  | Aztreonam BMD<sup>c,d</sup> |  | FDC+<br>4 µg/ml ATM | MIC fold<br>change<sup>e</sup> | Source |
| --- | --- | --- | --- | --- | --- | --- | --- | --- | --- | --- | --- |
|  |  |  |  |  | FDC MIC | Int. | ATM MIC | Int. |  |  |  |
| Ecc1 | *Enterobacter hormaechei* | ST2247* | NDM-1 | CTX-M-15 | 1 | S | 32 | R | 0.5 | 2 | this study (SAMN38656875) |
| Ecc2 | *Enterobacter hormaechei* | ST2713* | NDM-1 | CTX-M-15 | 1 | S | 64 | R | 1 | 1 | this study (SAMN38656876) |
| Ecc4 | *Enterobacter hormaechei* | ST171 | NDM-1 | CTX-M-15 | 4 | R | 64 | R | 4 | 1 | this study (SAMN38656877) |
| Ecc5 | *Enterobacter hormaechei* | ST171 | NDM-1 | CTX-M-15 | 4 | R | 64 | R | 2 | 2 | this study (SAMN38656878) |
| Ecc6 | *Enterobacter hormaechei* | ST270 | NDM-1 | - | 8 | R | 128 | R | 2 | 4 | this study (SAMN38656879) |
| Ecc7 | *Enterobacter hormaechei* | ST306 | NDM-1 | - | 2 | S | 0.125 | S | <0.0625 | 32 | this study (SAMN38656880) |
| Ecc8 | *Enterobacter hormaechei* | ST234 | NDM-1 | CTX-M-15 | 0.25 | S | 32 | R | 0.125 | 2 | this study (SAMN38656881) |
| Ecc9 | *Enterobacter hormaechei* | ST1439* | NDM-1 | CTX-M-15 | 1 | S | 8 | R | 0.5 | 2 | this study (SAMN38656882) |
| Ecc10 | *Enterobacter hormaechei* | ST1053 | NDM-1 | - | 1 | S | 4 | I | <0.0625 | 16 | this study (SAMN38656883) |
| Ecc11 | *Enterobacter hormaechei* | ST171 | NDM-1 | CTX-M-15 | 8 | R | 128 | R | 8 | 1 | this study (SAMN38656884) |
| Ecc12 | *Enterobacter hormaechei* | ST114 | NDM-1 | - | 1 | S | 128 | R | 1 | 1 | this study (SAMN38656885) |
| Ecc13 | *Enterobacter chuandaensis* | ST573* | NDM-5 | - | 16 | S | 0.25 | S | <0.0625 | 256 | this study (SAMN38656886) |
| Ecc14 | *Enterobacter hormaechei* | ST1439* | NDM-1 | CTX-M-15 | 1 | S | 16 | R | 1 | 1 | this study (SAMN38656887) |
| Ecc15 | *Enterobacter hormaechei* | ST48* | NDM-1 | - | 0.25 | S | 0.125 | S | <0.0625 | 4 | this study (SAMN38656888) |
| Ecc16 | *Enterobacter hormaechei* | ST953* | NDM-1 | CTX-M-15 | 2 | S | 32 | R | 2 | 1 | this study (SAMN38656889) |
| Ecc17 | *Enterobacter hormaechei* | ST51 | NDM-5 | - | 16 | R | 0.125 | S | <0.0625 | 256 | this study (SAMN38656890) |
| Ecc18 | *Enterobacter hormaechei* | ST306 | NDM-1 | - | 1 | S | 0.125 | S | <0.0625 | 16 | this study (SAMN38656891) |
| Ecc19 | *Enterobacter hormaechei* | ST90 | NDM-1 | - | 2 | S | 32 | R | 2 | 1 | this study (SAMN38656892) |
| Ecc20 | *Enterobacter hormaechei* | ST234 | NDM-1 | CTX-M-15 | 0.125 | S | 8 | R | <0.0625 | 2 | this study (SAMN38656893) |
| Ecc21 | *Enterobacter hormaechei* | ST1439* | NDM-1 | CTX-M-15 | 4 | R | 16 | R | 1 | 4 | this study (SAMN38656894) |
| etcl_1 | *Enterobacter hormaechei* | ST96 | NDM-5, OXA-48 | CTX-M-9 | 8 | R | 16 | R | 0.5 | 16 | SAMN19988671 [7] |
| etcl_1_mut | *Enterobacter hormaechei* | ST96 | NDM-5, OXA-48 | CTX-M-9 | >16 | R | 16 | R | >16 | 1 | SAMN20428006 [2] |

Abbreviations: MLST=multi-locus sequence type; NDM=New Delhi metallo-β-lactamase; OXA-48=oxacillinase 48; FDC=cefiderocol, ATM=aztreonam; BMD=broth microdilution; Int.=interpretation
<sup>a</sup> \* indicate the nearest MLST, derived from the whole-genome sequence data
<sup>b</sup> All clinical isolates from this study harboured the *bla*<sub>CTX-M-15</sub> gene. Isolate etcl\_1 harboured *bla*<sub>CTX-M-9</sub>.
<sup>c</sup> Antibiotic susceptibility testing interpretation based on EUCAST clinical breakpoints (v13.1); S=susceptible and R=resistant
<sup>d</sup> BMD was performed with iron-depleted cation-adjusted Mueller-Hinton broth, except for aztreonam only
<sup>e</sup> The MIC fold change is calculated by dividing cefiderocol MIC by cefiderocol-aztreonam MIC

Previously, we demonstrated that an NDM-producing *E. hormaechei*,etcl_1, was able to rapidly acquire cefiderocol resistance via mutation of the siderophore receptor CirA upon cefiderocol exposure, both *in vivo* and *in vitro*, and that the inhibition of the NDM-activity using dipicolinic acid was able to reduce the cefiderocol MIC and at the same time prevent the emergence of resistant clones [2]. Since the addition of 4 µg/ml aztreonam was able to reduce the cefiderocol MIC by 16-fold, we challenged the hypothesis that this MIC reduction could prevent the development/emergence of resistant mutants *in vitro* in a similar manner to dipicolinic acid.

In the initial resistance induction experiment, the development of cefiderocol resistance was observed for an inoculum size >10^5^ and >10^6^ cfu/ml at 4x MIC (32µg/ml) and 10x MIC (80µg/ml), respectively, when cefiderocol was used alone (Figure 1). Combining cefiderocol with aztreonam at a fixed concentration of 2 µg/ml did not affect the resistance development. However, when cefiderocol was combined with aztreonam at a fixed concentration of 4 µg/ml and 8 µg/ml, resistance development was only observed for an inoculum size of 10^8^ cfu/mL, indicating that the combination of cefiderocol-aztreonam could prevent the emergence of cefiderocol resistance *in vitro*.

**Figure 1.**
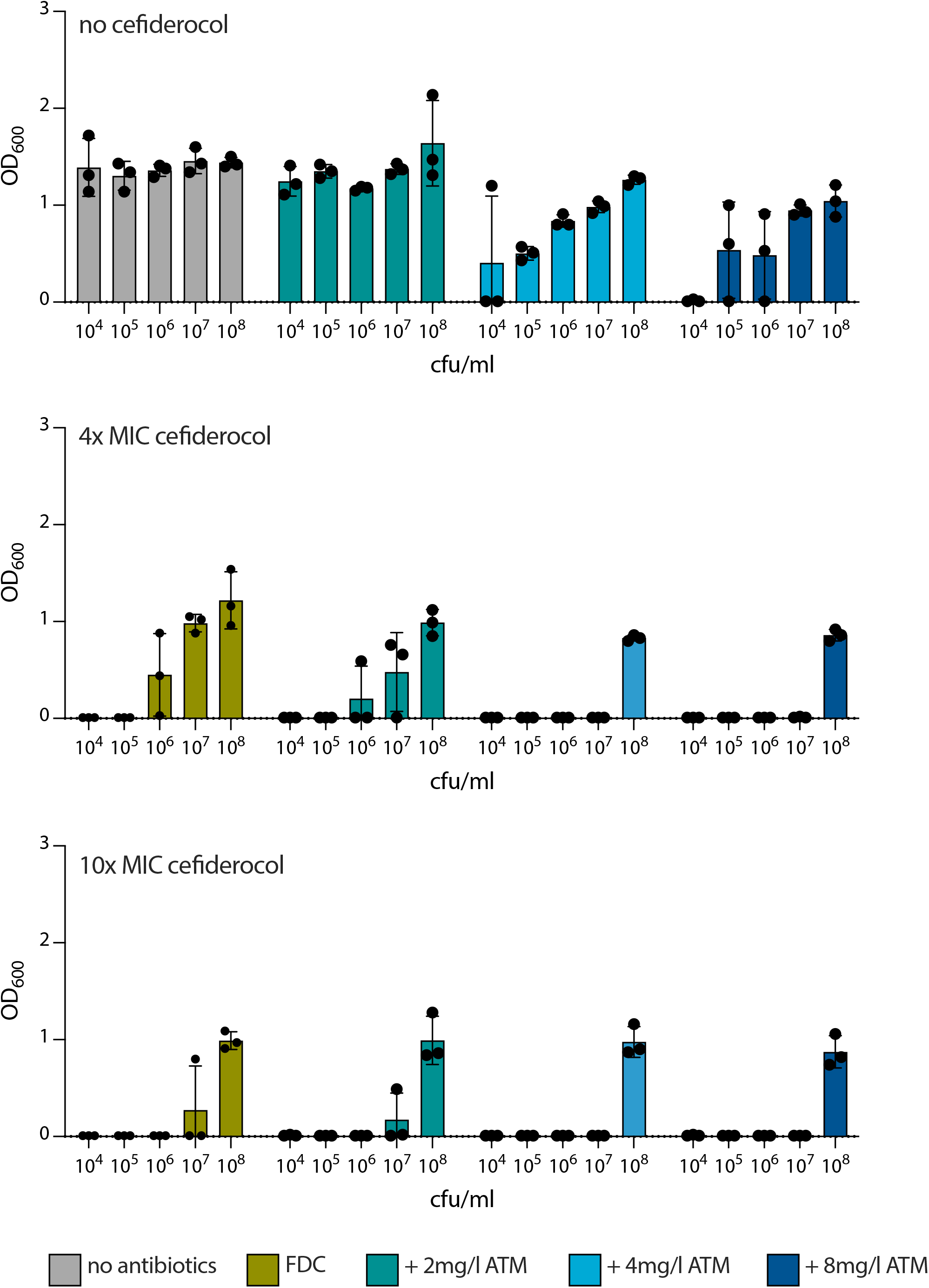
Resistance induction experiments. **(a)** antimicrobial activity of aztreonam using initial inoculums ranging from 10^4^ to 10^8^ cfu/ml. Aztreonam alone could not fully inhibit bacterial growth at concentrations of 2, 4, or 8 µg/ml. **(b)** Cefiderocol at 4x MIC could not fully inhibit bacterial growth at inoculum size ≥10^6^ cfu/ml. However, the addition of ≥4 µg/ml aztreonam was able to inhibit bacterial growth in three independent biological replicates consistently, indicating the absence of resistant mutants or subpopulations. **(c)** Similar to the findings for 4x MIC, 10x MIC inhibited the emergence and selection of resistance in vitro. At an inoculum of 10^8^ cfu/ml, none of the tested substances could fully inhibit bacterial growth. All experiments were performed as biological triplicates. Bacterial growth was quantified as optical density at 600nm (OD600). Abbreviations: FDC=cefiderocol, ATM=aztreonam, MIC=minimal inhibitory concentration.

**Figure 2.**
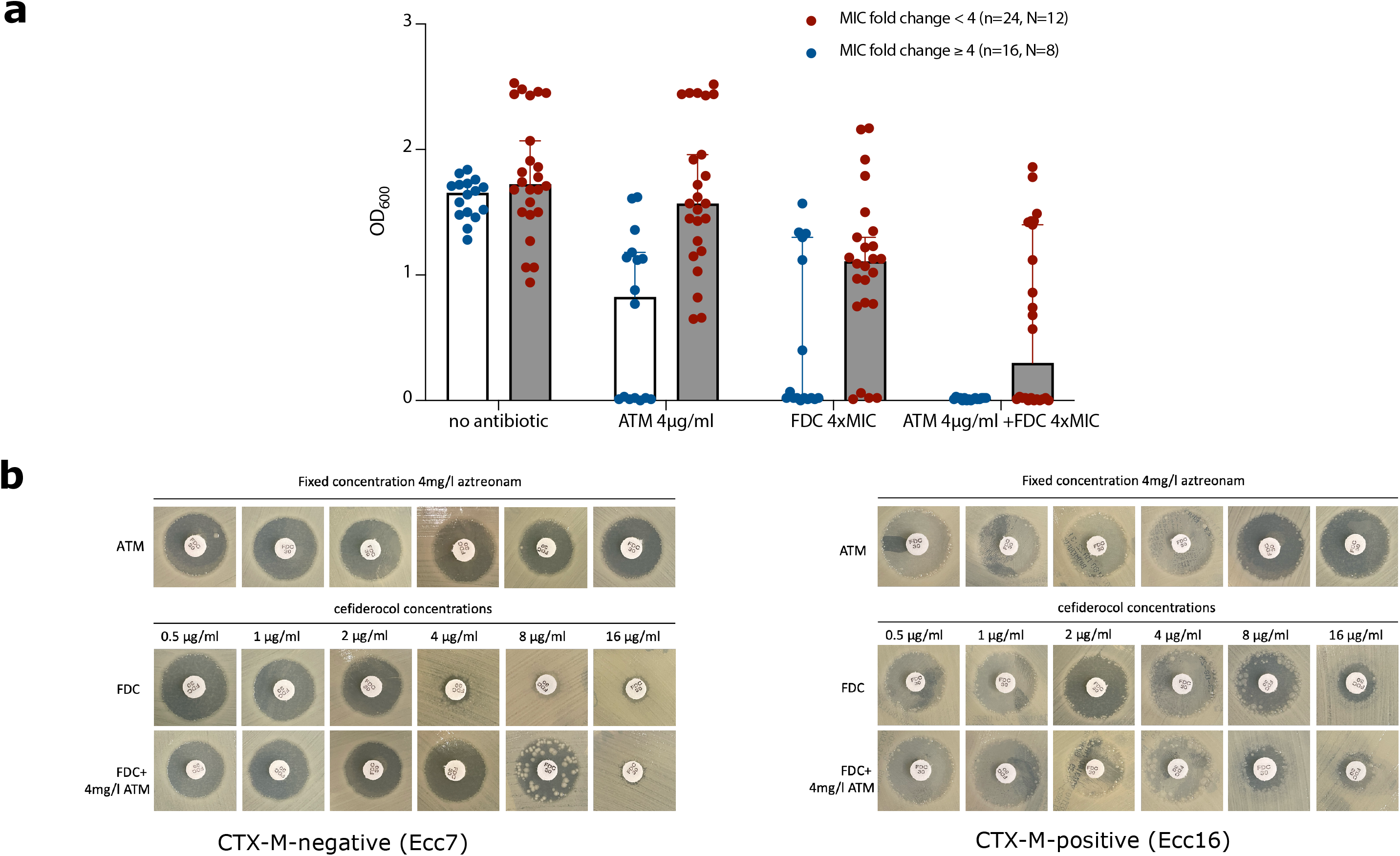
Validation of resistance induction experiments using clinical isolates. **(a)**. The resistance induction experiment at 4x cefiderocol (FDC) MIC with or without a fixed concentration of 4 µg/ml aztreonam (ATM) using 20 NDM-producing *E. cloacae* complex clinical isolates. Two biological replicates (n) are plotted for each isolate (N). Combining cefiderocol and 4µg/ml aztreonam inhibited all isolates (blue circles) with a MIC fold change ≥4, but not isolates with MIC fold change <4 (red circles). **(b)** Serial passaging experiment with doubling concentration per passage of isolate Ecc7 (CTX-M-15 negative, MIC fold change 32) and Ecc16 (CTX-M-15 positve, MIC fold change 1). Combining cefiderocol with a fixed concentration of 4 µg/ml aztreonam was able to delay the emergence of cefiderocol-resistant clones.

### Validation of initial observation using NDM-producing E. cloacae complex clinical isolates

Next, we validated this finding using randomly chosen 20 clinical isolates of NDM-producing *E. cloacae* complex. The molecular characteristics and the phenotypic resistance towards cefiderocol, aztreonam and cefiderocol-aztreonam were summarized in Table 1. Of 20 isolates, 8 (40%) exhibited a MIC reduction (ΔMIC) of ≥4-fold, which is generally considered a significant reduction. Indeed, in the resistance induction experiment using an inoculum of 10^7^ cfu/mL, we observed that resistance development can be inhibited consistently in two independent biological replicates for all 8 of 20 isolates with with a ΔMIC ≥4-fold (Supplementary Figure S3a). Next, we performed a serial passaging experiment by doubling the cefiderocol concentrations in each passage while keeping the aztreonam constant at 4 µg/ml throughout for the isolates Ecc7 (ΔMIC=32-fold) and Ecc16 (ΔMIC=no change). For Ecc7, the resistance clones appeared after the cefiderocol concentration reached 4µg/ml. Whereas in the combination cefiderocol-aztreonam, some resistant clones appeared only after reaching a cefiderocol concentration of 8µg/ml. In contrast, for Ecc16, resistance clones emerged more rapidly in the combination cefiderocol-aztreonam than cefiderocol alone. For both isolates, aztreonam alone did not induce resistance towards cefiderocol.

Phenotypic susceptibility towards aztreonam is not a good predictor for a significant ΔMIC. Although all 12 isolates (12/20, 60%) with a non-significant ΔMIC were all phenotypically aztreonam resistant, two of the six isolates with a significant ΔMIC were resistant to aztreonam (8 to 2 µg/ml for Ecc6 and 4 to 1 µg/ml for Ecc21, Table 1).

To identify potential genomic markers, we performed WGS on all 20 isolates. Unsurprisingly, isolates resistant to cefiderocol did not exhibit mutations of the CirA genes as previously described, as this type of mutation typically confers a high MIC (>256 µg/ml) [7, 8]. We also did not observe a phylogenetic correlation with the reduction of the cefiderocol MIC, as this reduction occurs in different clones and strains. Interestingly, although AmpC is chromosomally encoded in Enterobacter species, the presence of AmpC-encoding genes was not associated with aztreonam susceptibility or ΔMIC (Supplementary Figure S3a). Using pangenome-wide genomic analysis [9], we identified the significant positive association (p<0.001) between significant ΔMIC and genes which were co-localized, therefore showing identical effect size and standard deviation, amongst which *bla*_OXA-1_ and *bla*_CTX-M-15_ (Supplementary Figure S3b). However, our *in vitro* experimental data suggest that the best predictor for the inhibition of resistance development was a ΔMIC of 4-fold or more.

## Discussion

Rapid emergence and selection of cefiderocol resistance during therapy have been reported in various NDM-producing species, suggesting a low threshold for resistance development associated with NDM production [2, 7, 8]. Indeed, NDM-type β-lactamases have been demonstrated to have slight hydrolytic activity towards cefiderocol, which may explain the elevated cefiderocol MIC and high propensity to develop cefiderocol resistance in NDM-producers [10, 11]. While these *in vivo* observations can and have been replicated in experimental *in vitro* settings numerously, studies exploring the possibilities to prevent the emergence and selection of cefiderocol resistance are lacking.

Our *in vitro* study demonstrated that by combining cefiderocol with aztreonam, the emergence and selection of resistant mutants can be prevented/delayed in NDM-producing clinical *E. cloacae* complex isolates for inoculum size less than 10^8^ cfu/ml. Consistent with the literature, the inoculum size had a significant impact on the MIC of both β-lactam antibiotics and the emergence of resistant mutants [12]. By performing validation of our initial finding using 20 clinical NDM-producing *E. cloacae* complex isolates, we could demonstrate that a MIC change of at least 4-fold is a good predictor for the inhibition of resistance development *in vitro*. Although some NDM-producing isolates were phenotypically resistant to aztreonam (MIC ≥8 µg/ml), a lower aztreonam concentration of 4 µg/ml combined with cefiderocol was sufficient to prevent the emergence of resistance in the tested isolates. Based on pharmacokinetic and pharmacodynamic data on aztreonam, with a c_max_ of >100µg/ml following i.v. dosing of 1 or 2 g [13] and plasma concentrations of >10µg/ml 1 hour post-infusion [14, 15], it is plausible to assume that the aztreonam concentration used in our *in vitro* experiments is within the expected *in vivo* therapeutic concentration range.

Although some of the tested *E. cloacae* complex isolates were phenotypically resistant towards cefiderocol (elevated MIC), the combination cefiderocol-aztreonam at a fixed aztreonam concentration of 4 µg/ml could reduce the MIC to a susceptible range based on the EUCAST clinical breakpoint for cefiderocol only (≤4 µg/ml). Thus, supporting the notion that elevated cefiderocol MIC is associated with rapid emergence and selection of cefiderocol resistance [2, 7]. Meanwhile, combining cefiderocol with aztreonam does not affect the MIC for cefiderocol-resistant *E. cloacae* complex due to the acquisition of mutations in the catecholate siderophore receptor CirA. In *E. hormaechei* etcl_1_mut isolate with CirA frameshift mutations (G2710T), cefiderocol/aztreonam combination did not reduce the cefiderocol MIC (≥16 µg/ml), confirming that CirA-mutation-mediated resistance phenotype is independent of NDM-production and suggesting that this resistance mechanism cannot be overcome by combining cefiderocol with aztreonam.

The idea of combining cefiderocol with another substance to increase the efficacy of this antibiotic substance is not entirely novel. Some studies have explored the microbiological effects of combining cefiderocol with novel β-lactam/β-lactamase inhibitors, such as ceftazidime/avibactam or β-lactamase inhibitors alone, such as tazobactam, avibactam, vaborbactam, and relebactam [16, 17]. However, these studies only studied the antibacterial effect of such combinations and did not study the effect of various cefiderocol combinations on the propensity of resistance emergence and selection. To the best of our knowledge, the combination of cefiderocol/aztreonam has not been studied extensively so far.

Combining two substances of the same substance class, e.g. dual β-lactam therapy, may be counterintuitive, but this combination may be useful in inactivating multiple β-lactamases simultaneously to achieve synergistic bacterial killing and minimize the risk of resistance induction [18]. Some bacterial species, such as *Enterobacter* species harbour intrinsic β-lactamases, such as AmpC, and sometimes even additional β-lactamases, such as CTX-M, OXA-48 and NDM. In the case of cefiderocol/aztreonam, cefiderocol is stable against weaker β-lactamases but can be slightly hydrolyzed by NDM. Meanwhile, aztreonam is stable against NDM but can be hydrolyzed by weaker β-lactamases, such as CTX-M, AmpC and OXA-48 [19]. Therefore, by combining the two substances, a cross-protection, meaning that cefiderocol is protecting aztreonam hydrolysis by NDM and vice versa, rather than synergy may explain the MIC reduction compared to the mono substances. Moreover, the sequential exposure of β-lactam antibiotics may promote the microbiological efficacy of β-lactams and increase the threshold for resistance development. Our data suggest that despite intrinsic AmpC in *Enterobacter* spp, aztreonam alone or combined with cefiderocol retains antimicrobial activity. However, the presence of *bla*_CTX-M_ plasmid was associated with failure in preventing the emergence of cefiderocol-resistant clones using the combination cefiderocol-aztreonam. Although double β-lactam combinations are not yet common practice for treating infections due to Enterobacterales, a double β-lactam combination (ampicillin and ceftriaxone) has been explored for *Enterococcus faecalis* and is now recommended for the treatment of *E. faecalis* bloodstream infections and infective endocarditis [20].

Our study has limitations. Our data and observation are based on *in vitro* observations, and *in vivo* validation is still needed. Further, we have only focused on *E. cloacae* complex, which limits the generalizability of our findings. The effect of cefiderocol-aztreonam on other Enterobacterales is unknown and warrant further investigations. Despite these limitations, we have validated and were able to replicate our initial findings using 20 NDM-producing clinical strains. Considering the cumulating reports on therapy-related development of cefiderocol resistance, our *in vitro* data gives insight into the possibilities of combining cefiderocol with aztreonam to prevent cefiderocol resistance emergence in NDM-producing *E. cloacae* complex. The combination of dual β-lactam antibiotics to overcome and inactivate multiple β-lactamases may be an interesting and valuable strategy for preventing the rapid development of resistance to novel β-lactam antibiotics in multidrug-resistant bacteria, such as carbapenemase-producing Enterobacterales.

## Author contributions

Study concept: LG, SB, DN Experiments: LG, TNMT, ÖC, TTT Bioinformatics analysis: SB Statistical analysis and visualization: SB, DN Supervision: JR, DN Drafting: LG, SB, SH, JR, DN Finalisation of manuscript: all authors

## Acknowledgements

The authors thank Melanie Albrecht for her excellent technical assistance.

## Data availability

Draft genomes are available in the Bioproject PRJNA1048675.

## References

[1] Yamano Y. In Vitro Activity of Cefiderocol Against a Broad Range of Clinically Important Gram-negative Bacteria. Clin Infect Dis. 2019;69:S544–S51.

[2] Nurjadi D, Kocer K, Chanthalangsy Q, Klein S, Heeg K, Boutin S. New Delhi Metallo-Beta-Lactamase Facilitates the Emergence of Cefiderocol Resistance in Enterobacter cloacae. Antimicrob Agents Chemother. 2022;66:e0201121.

[3] Falcone M, Daikos GL, Tiseo G, Bassoulis D, Giordano C, Galfo V, et al. Efficacy of Ceftazidime-avibactam Plus Aztreonam in Patients With Bloodstream Infections Caused by Metallo-beta-lactamase-Producing Enterobacterales. Clin Infect Dis. 2021;72:1871–8.

[4] Tamma PD, Aitken SL, Bonomo RA, Mathers AJ, van Duin D, Clancy CJ. Infectious Diseases Society of America 2022 Guidance on the Treatment of Extended-Spectrum beta-lactamase Producing Enterobacterales (ESBL-E), Carbapenem-Resistant Enterobacterales (CRE), and Pseudomonas aeruginosa with Difficult-to-Treat Resistance (DTR-P. aeruginosa). Clin Infect Dis. 2022;75:187–212.

[5] Sandfort M, Hans JB, Fischer MA, Reichert F, Cremanns M, Eisfeld J, et al. Increase in NDM-1 and NDM-1/OXA-48-producing Klebsiella pneumoniae in Germany associated with the war in Ukraine, 2022. Euro Surveill. 2022;27.

[6] Hackel MA, Tsuji M, Yamano Y, Echols R, Karlowsky JA, Sahm DF. Reproducibility of broth microdilution MICs for the novel siderophore cephalosporin, cefiderocol, determined using iron-depleted cation-adjusted Mueller-Hinton broth. Diagn Microbiol Infect Dis. 2019;94:321–5.

[7] Klein S, Boutin S, Kocer K, Fiedler MO, Storzinger D, Weigand MA, et al. Rapid Development of Cefiderocol Resistance in Carbapenem-resistant Enterobacter cloacae During Therapy Is Associated With Heterogeneous Mutations in the Catecholate Siderophore Receptor cirA. Clin Infect Dis. 2022;74:905–8.

[8] McElheny CL, Fowler EL, Iovleva A, Shields RK, Doi Y. In Vitro Evolution of Cefiderocol Resistance in an NDM-Producing Klebsiella pneumoniae Due to Functional Loss of CirA. Microbiol Spectr. 2021;9:e0177921.

[9] Lees JA, Galardini M, Bentley SD, Weiser JN, Corander J. pyseer: a comprehensive tool for microbial pangenome-wide association studies. Bioinformatics. 2018;34:4310–2.

[10] Poirel L, Sadek M, Nordmann P. Contribution of PER-Type and NDM-Type beta-Lactamases to Cefiderocol Resistance in Acinetobacter baumannii. Antimicrob Agents Chemother. 2021;65:e0087721.

[11] Ito-Horiyama T, Ishii Y, Ito A, Sato T, Nakamura R, Fukuhara N, et al. Stability of Novel Siderophore Cephalosporin S-649266 against Clinically Relevant Carbapenemases. Antimicrob Agents Chemother. 2016;60:4384–6.

[12] Hobson CA, Cointe A, Jacquier H, Choudhury A, Magnan M, Courroux C, et al. Cross-resistance to cefiderocol and ceftazidime-avibactam in KPC beta-lactamase mutants and the inoculum effect. Clin Microbiol Infect. 2021;27:1172 e7-e10.

[13] Smith PF, Ballow CH, Booker BM, Forrest A, Schentag JJ. Pharmacokinetics and pharmacodynamics of aztreonam and tobramycin in hospitalized patients. Clin Ther. 2001;23:1231–44.

[14] Swabb EA. Review of the clinical pharmacology of the monobactam antibiotic aztreonam. Am J Med. 1985;78:11–8.

[15] Ramsey C, MacGowan AP. A review of the pharmacokinetics and pharmacodynamics of aztreonam. J Antimicrob Chemother. 2016;71:2704–12.

[16] Boattini M, Comini S, Bianco G, Iannaccone M, Casale R, Cavallo R, Costa C. Activity of cefiderocol and synergy of novel beta-lactam-beta-lactamase inhibitor-based combinations against metallo-beta-lactamase-producing gram-negative bacilli: insights from a two-year study (2019-2020). J Chemother. 2023;35:198–204.

[17] Palombo M, Bovo F, Amadesi S, Gaibani P. Synergistic Activity of Cefiderocol in Combination with Piperacillin-Tazobactam, Fosfomycin, Ampicillin-Sulbactam, Imipenem-Relebactam and Ceftazidime-Avibactam against Carbapenem-Resistant Gram-Negative Bacteria. Antibiotics (Basel). 2023;12.

[18] Jiao Y, Moya B, Chen MJ, Zavascki AP, Tsai H, Tao X, et al. Comparable Efficacy and Better Safety of Double beta-Lactam Combination Therapy versus betaLLactam plus Aminoglycoside in Gram-Negative Bacteria in Randomized, Controlled Trials. Antimicrob Agents Chemother. 2019;63.

[19] Sy SK, Beaudoin ME, Zhuang L, Loblein KI, Lux C, Kissel M, et al. In vitro pharmacokinetics/pharmacodynamics of the combination of avibactam and aztreonam against MDR organisms. J Antimicrob Chemother. 2016;71:1866–80.

[20] Beganovic M, Luther MK, Rice LB, Arias CA, Rybak MJ, LaPlante KL. A Review of Combination Antimicrobial Therapy for Enterococcus faecalis Bloodstream Infections and Infective Endocarditis. Clin Infect Dis. 2018;67:303–9.

