## Supplementary methods and figures for "Delaying cefiderocol resistance development in NDM-producing *Enterobacter cloacae* complex by combining cefiderocol with aztreonam *in vitro*"

**Title:**

**Supplementary Method**

*Checkerboard assay to test for synergy.*

To determine the synergistic effect of cefiderocol and aztreonam, a modified checkerboard assay was used [1]. Briefly, a 96-well microplate with concentrations of cefiderocol and aztreonam ranging from 0.5 µg/mL to 32 µg/mL was prepared with fresh iron-depleted CA-MHB. For a final inoculum concentration in the microplate of 5 x 10^5^ CFU/mL, 0.5 McFarland standard (comparable to a bacterial suspension of 1.5 x 10^8^ CFU/mL) was adjusted in fresh iron-depleted CA-MHB and diluted 1:100 in fresh iron-depleted CA-MHB. The microplate was incubated at 37°C and the optical density at 600nm (OD_600_) was measured after 18±2 h using an Infinite M200 PRO (Tecan, Switzerland). The following formula was used for the calculation of fractional inhibitory concentration index (FICI)= (MIC_substance_ _A_ in combination/MIC_substance A_ alone) + (MIC_substance B_ in combination/MIC_substance B_ alone). The effects of the antibiotic agent combinations were classified according to the following criteria: (1) FICI ≤ 0.5, synergistic effects; (2) 0.5 < FICI ≤ 1, additive effects; (3) 1 < FICI < 4, no interactions; (4) FICI ≥ 4.0, antagonistic effects.

*Genome sequencing and bioinformatics analysis*

DNA extraction was performed from fresh culture on BD™ Columbia Agar with 5% Sheep Blood (Becton Dickinson GmbH, Heidelberg, Germany) at 37 °C. DNA was extracted using the DNeasy Blood and Tissue Kit (Qiagen GmbH, Hilden, Germany) according to the manufacturer’s protocol. Library preparation was performed using Nextera DNA Flex Library Prep Kit (Illumina) and sequencing was done on a MiSeq Illumina platform (short-read sequencing, 2 ×301 bp). Post-sequencing procedure was performed as follows: Raw sequences were controlled for quality and adapter removal using fastp (v0·23·2 with parameters -q = 30 and -l = 45). Clean reads were then used to create de novo assembly using SPAdes 3.15.5 (with the option —careful and—only-assembler)[2, 3]. Draft genomes were curated by removing contigs with a length <500 bp and/or coverage <10×. The quality of the final draft was quality-controlled using Quast (v5.0.2) [4]. The complete draft genomes were processed through available databases using Abricate (minimum identity 90% and minimum coverage 80%) (<https://github.com/tseemann/abricate>) to identify antimicrobial resistance (NCBI, CARD, ARG-ANNOT, ResFinder, MEGARES databases), virulence genes (VFDB databases) and plasmid type (PlasmidFinder database) to identify the Inc type of the plasmid. Each genome was annotated using pgap (version 2023-10-03.build7061). The core genome (Prevalence = 100%) was assessed using Roary (v 3.13.0) and the alignment was then corrected with Gubbins 3.2.1 to create a recombination-corrected phylogeny. The gene presence/absence table from Roary was used to associate gene variability with the phenotypic resistance to the combination of aztreonam and cefiderocol using Pyseer v1.3.11 with a fixed model including the recombination adjusted phylogenetic distance and using 2 dimensions based on the MDS eigenvalues (Supplementary Figure S1). Only genes associated with a Likelihood ratio test p-value ≤ 0.01 were considered as significantly associated with the phenotype.

**References:**

[1] Bellio P, Fagnani L, Nazzicone L, Celenza G. New and simplified method for drug combination studies by checkerboard assay. MethodsX. 2021;8:101543.

[2] Bankevich A, Nurk S, Antipov D, Gurevich AA, Dvorkin M, Kulikov AS, et al. SPAdes: a new genome assembly algorithm and its applications to single-cell sequencing. J Comput Biol. 2012;19:455-77.

[3] Chen S, Zhou Y, Chen Y, Gu J. fastp: an ultra-fast all-in-one FASTQ preprocessor. Bioinformatics. 2018;34:i884-i90.

[4] Gurevich A, Saveliev V, Vyahhi N, Tesler G. QUAST: quality assessment tool for genome assemblies. Bioinformatics. 2013;29:1072-5.

**Supplementary figure S1.** Scree plot representing the eigenvalue of each dimension of the nonparametric multidimensional scaling (MDS) based on the phylogeny calculated by Gubbins. We could observe a drop after the second dimensions and therefore the first two dimensions only were user in the fixed model of the GWAS analysis.

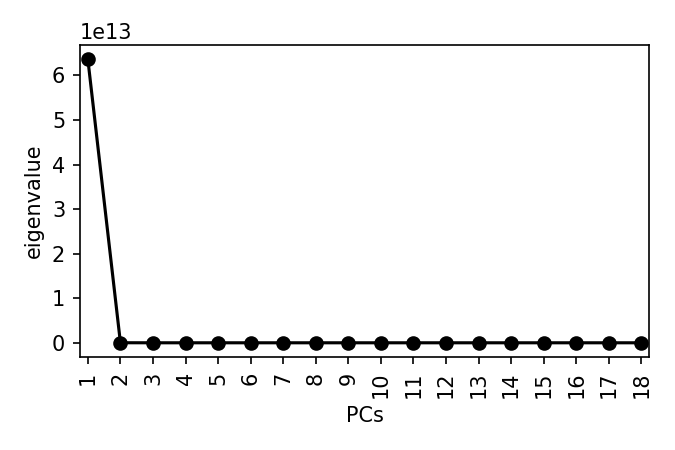

**Supplementary Figure S2. Synergy testing for cefiderocol and aztreonam. (a)** checkerboard assay for cefiderocol and aztreonam for *E. hormaechei* etcl_1. No synergistic effect was observed for the two tested substances. **(b)** Disk diffusion assay to test for synergy between cefiderocol and aztreonam. Placing the disk with a distance of 1 mm did not lead to distortions or enlargement of the zones of inhibitions, indicating that there was no synergistic effect for cefiderocol and aztreonam.

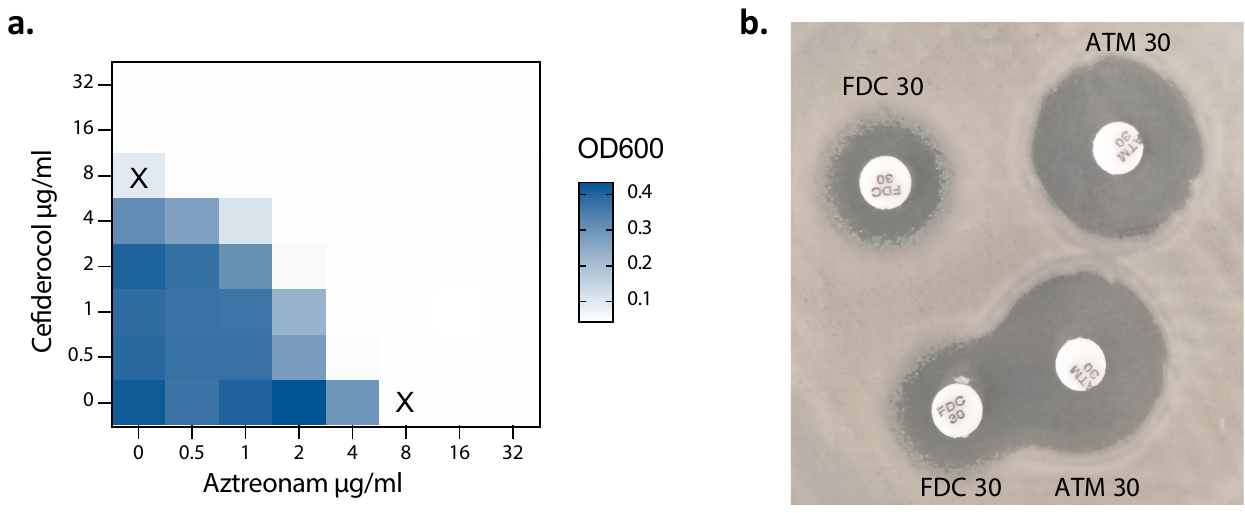

**Supplementary Figure S3. Genomic characteristics of clinical *Enterobacter cloacae* complex. (a)** Phylogenetic tree based on core genome SNP. The MIC fold change (MIC_FC) was calculated by dividing FDC+ATM MIC by FDC MIC. The aztreonam concentration was kept at a fixed concentration of 4 µg/ml. AST was interpreted using the EUCAST clinical breakpoints (v13.1). **(b)** GWAS analysis associating microbial pangenome presence/absence with the MIC fold change. The analysis was performed using Pyseer v1.3.11. The p-value is based on the likelihood ratio test and was considered significant if ≤ 0.01. A positive effect size indicated that the gene presence is associated with a higher MIC fold change. Abbreviations; MIC=minimum inhibitory concentration, ATM=aztreonam, FDC=cefiderocol, FC=fold change, AST=antibiotic susceptibility testing.
